# Persistent *Salmonella* infections in humans are associated with mutations in the BarA/SirA regulatory pathway

**DOI:** 10.1101/2022.06.21.496976

**Authors:** Alexandra Grote, Bar Piscon, Abigail L. Manson, Jonathan Livny, Ashlee M. Earl, Ohad Gal-Mor

## Abstract

The bacterial pathogen *Salmonella enterica* is able to establish persistent infections, evading antibiotics and the host immune system and providing a reservoir for recrudescence and transmission to new hosts. Non-typhoidal serovars (NTS) of *S. enterica* can establish and maintain symptomatic and asymptomatic long term human infections that may predispose carriers to inflammatory bowel diseases and cancer. Defining the adaptations and host-pathogen interactions enabling these persistent infections is key to devising more effective strategies to combat and prevent persistent bacterial infections. Using comparative genomics of 639 S*almonella* NTS isolates belonging to 49 serovars that were longitudinally obtained from 256 salmonelosis patients during different stages of infection, we identified numerous genetic variations accruing over time in strains isolated from the same patient. Many of these changes were found in the same gene across multiple patients and serovars. Among these variant loci, genes encoding global transcriptional regulators were found to be the most commonly mutated between early and late same-patient isolates. Genetic changes in the SirA/BarA two-component regulatory system were particularly frequent, with mutations identified in 24 independent patients. Comparative RNA-Seq analysis revealed that distinct mutations in *sirA/barA* that arose independently in late isolates of multiple patients lead to significantly diminished expression of virulence-associated genes encoded in the *Salmonella* Pathogenicity Islands (SPIs) 1 and 4, many of which are known to be critical for host cell invasion and the production of enteritis. Using the salmonellosis mouse model we showed that these mutations in *sirA/barA* genes confer attenuated virulence *in-vivo*. Taken together, these data suggest that selection of mutations in the SirA/BarA pathway facilitates persistent *Salmonella* infection in humans, possibly by attenuating *Salmonella* virulence and its ability to cause inflammation.

## INTRODUCTION

Different clinically-important bacterial pathogens, including *Mycobacterium tuberculosis*, *Helicobacter pylori*, *Pseudomonas aeruginosa,* and *Salmonella enterica,* are able to establish persistent infections in humans. These persistent bacteria can evade the host immune system and killing by antibiotics and have been linked to the emergence of new antibiotic resistant strains (N. R. Cohen et al., 2013). *S. enterica* is a gram-negative, facultative intracellular pathogen with over 2,600 antigenically-distinct serovars (Issenhuth-Jeanjean et al., 2014), the majority of which are considered non-typhoidal serovars (NTS) and frequently colonize food-producing and wild animals. Human infection by *S. enterica* serovars may cause different clinical outcomes, including asymptomatic colonization, gastroenteritis in the terminal ileum and colon, invasive systemic disease and bacteremia, and enteric (typhoid) fever. The precise outcome of a *Salmonella* infection is dependent on the characteristics of the infecting serovar and the immunological status of the infected individual (Gal-Mor, 2019). *S. enterica* infections still pose a significant global clinical challenge with over 27 million cases of enteric fever (Crump et al., 2004) and 78.7 million cases of gastroenteritis annually (Havelaar et al., 2015).

The two main hallmarks of *S. enterica* pathogenicity are the ability to invade nonphagocytic cells and to replicate within phagocytic and nonphagocytic host cells. These phenotypes are dependent on the function of two separate type three secretion systems (T3SSs) and their designated secreted effector proteins. T3SS-1 is encoded on *Salmonella* pathogenicity island (SPI)-1, expressed when *Salmonella* are extracellular and mediates its active invasion into nonphagocytic cells (Galán, 2001; Mills et al., 2006). Additionally, the products of SPI-1 genes are responsible for regulation of the host immune response (Pavlova et al., 2011; Zhao et al., 2018), and induction of neutrophil recruitment during enteric colitis, which lead to reduction and alteration of the intestinal microbiota (Sekirov et al., 2010; Winter et al., 2010). In contrast, T3SS-2, encoded on SPI-2, is induced when *Salmonella* is intracellular and is required for its intracellular survival and replication (Ochman et al., 1996; Shea et al., 1996).

SPI-1 gene regulation is complex and involves multiple regulators, encoded both within and outside of SPI-1 (Lou et al., 2019) that control gene expression at the transcriptional and post-transcriptional levels (Lou et al., 2019). The SirA/BarA two-component regulatory system (TCRS) is positioned at the top of the regulatory pathway, consisting of BarA, the sensor kinase, and SirA, the response regulator (Teplitski et al., 2003). The SirA/BarA system can activate the SPI-1 transcriptional regulators HilA, HilC, HilD and InvF (Behlau & Miller, 1993; Ellermeier & Slauch, 2007; Teplitski et al., 2003). In addition, SirA/BarA system relays SPI-1 gene downregulation by activating the expression of *csrB/C*, two non-coding regulatory RNAs. *csrB/C* sequester the RNA-binding protein CsrA, inhibiting CsrA from binding to the *hilD* mRNA and preventing HilD translation (Martínez et al., 2011; Teplitski et al., 2003). This SirA/BarA TCRS is well-conserved across different bacteria and has orthologs in *Pseudomonas* species (GagA/GacS), *Vibrio cholerae* (VarA/VarS), *Erwinia carotovora* (ExpA/ExpS), *Legionella pneumophila* (LetA/LetS), and *Escherichia coli* (UvrY/BarA) where it is similarly involved in regulation of virulence, antibiotic production, motility, and biofilm production (Heeb & Haas, 2001).

While it is well-known that typhoidal serovars cause persistent salmonellosis (Foster et al., 2021; Gal-Mor, 2019; J Barton et al., 2021), we previously showed that NTS can also establish symptomatic or asymptomatic persistent infections in humans at a frequency of at least 2.2%, which can last months to years (Marzel et al., 2016). While the underlying mechanisms of *Salmonella* persistence are poorly understood, studies of NTS infections in humans and mouse models have shed some light on host, bacterial and environmental factors that contribute to persistent salmonellosis, including the effect of gender and age (Buchwald & Blaser, 1984; Dixon, 1965; Musher & Rubenstein, 1973; Vogelsang & Bøe, 1948), and antibiotic treatment (Stecher & Hardt, 2011) on long-term *Salmonella* infections. In our previous study of paired whole genome sequences of early and late isolates from 11 patients with *S*. Typhimurium infections, single nucleotide polymorphisms (SNPs) were found in global virulence regulatory genes, including *dksA*, *rpoS*, *hilD*, *melR*, *rfc*, and *barA* (Marzel et al., 2016). Nonetheless, no convergence at the variant or gene level was identified between patients. Similarly, in a study of whole genomes from an *S*. Typhimurium infection lasting 55 days, a nonsynonymous mutation was found in FlhC, a master regulator of flagellar biosynthesis (Octavia et al., 2015). While neither of these studies was able to identify conserved mutations across multiple patients, these singular examples point to a role for key virulence regulators in the development of persistent salmonelosis. Nevertheless, little is known about the molecular pathways that enable persistence or how this process evolves during human infections.

We therefore sought to expand our previous observations in order to find dominant routes through which *Salmonella* evolves to persist long-term in a human host. To this end we used comparative genomics of early and late isolates from 256 different patients drawn from a large retrospective study of *Salmonella* persistence including 48,345 culture-confirmed salmonellosis cases that occurred in Israel over a 17 year period between 1995 and 2012. Furthermore, we go on to characterize the transcriptional and phenotypic differences between early and late same-patient isolates with convergent mutations across multiple patients and demonstrate that these mutations lead to attenuated virulence *in vivo*.

## RESULTS

### Persistent infections are caused by a diverse set of *Salmonella* serovars

To profile adaptations leading to persistence across NTS, we collected and whole genome sequenced all *S. enterica* isolates obtained from patients in Israel presenting with persistent symptomatic salmonellosis between 1995 and 2012 (PRJNA847966) (Marzel et al., 2016) [**Supplemental Table 1**]. Each patient was represented by 2-5 independent isolates obtained during different stages of infection that lasted between 30 and 2,001 days from the first culture-confirmed diagnosis. Of the 639 isolates (obtained from 256 patients) sequenced as part of this study, analysis using Centrifuge (Kim et al., 2016) revealed that data from 8 samples (from 3 different patients) showed high amounts of non-*Salmonella* bacterial DNA, likely due to co-infection, and were therefore excluded from further analysis. Sequence reads from each of the remaining 631 isolates were aligned to the reference genome of *S*. Typhimurium SL1344 (NC_016810.1) to identify variants, including single nucleotide polymorphisms (SNPs), which were used as input into a maximum likelihood-based phylogenetic analysis to solve the evolutionary relationships among isolates. Mapping the results of laboratory serotyping onto the phylogeny revealed largely congruent results, with each of the 49 predicted serovars corresponding to a closely related cluster on the phylogenetic tree [**Figure 1A**].

**Figure 1:**
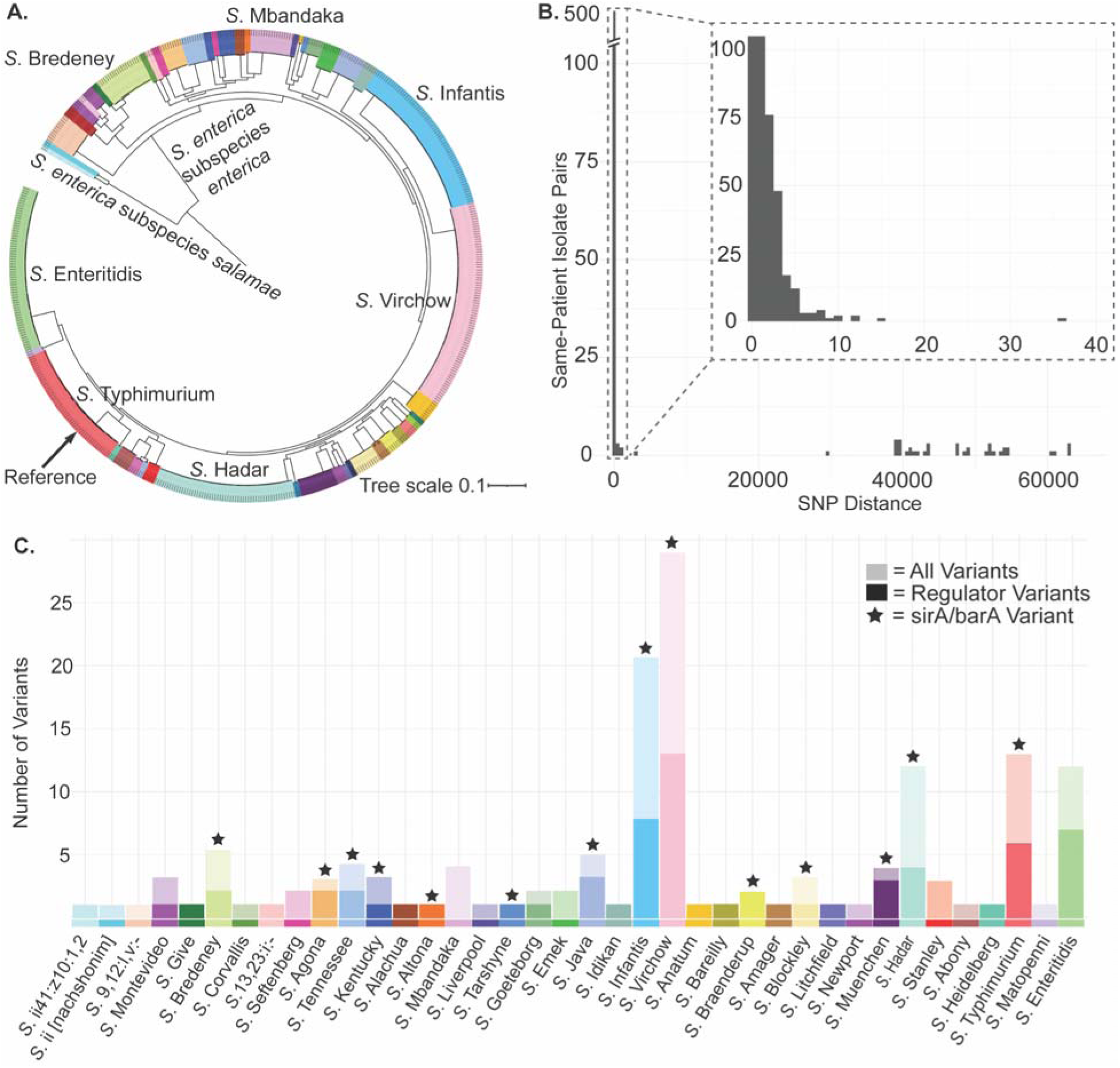
Genetic diversity of *S. enterica* isolates during persistent infection. A) Phylogeny of 562 *S. enterica* isolate genomes from human persistent infections. The colored ring represents serovar information as determined by serotyping according to the White-Kauffmann-Le Minor scheme (Issenhuth-Jeanjean et al., 2014). B) Distribution of SNP distances between 545 isolate pairs, obtained from the same patient at different time points. C) Variants, including SNPs, insertions and deletions, in all genes, and in genes with the GO term for “Regulation of Biological Process” (GO:0050789), between early and late same-patient isolates. Bars are colored by serovar, with light tone representing all variants and dark tone representing variants in genes with GO:0050789.

These analyses showed that persistent salmonelosis can be caused by many NTS serovars, however, in support of our previous work (Marzel et al., 2016), we found serovars Mbandaka, Bredeney, Infantis, and Virchow enriched in our dataset of persistent isolates compared to their general prevalence in “regular” sporadic clinical isolates, indicating their apparent propensity to cause persistent infections (Marzel et al., 2016).

### Most recurrent infections are due to persistence rather than reinfection

Though most same-patient isolates fell within the same serovar, we sought to differentiate cases of persistent infection by the same strain from re-infection by a similar strain. To do this, we performed pairwise comparisons of isolates from the same patient using variants called against the *S*. Typhimurium SL1344 reference genome, considering the first culture-confirmed isolate as the “early” isolate and the last culture-confirmed isolate as the “late” isolate. 88% (222 of 253) of patients had isolates separated by 36 SNPs or fewer based on the maximum length of infection in our study and previously published SNP rates (Marzel et al., 2016) [**Figure 1B**]. This threshold includes 72 patients (28%) with zero SNPs separating their isolates and 31 patients with isolates separated by more than 200 SNPs, likely representing reinfections or original non-clonal infections, rather than persistence by a single clone. The number of SNPs accumulating over time, on average, across pairs was found to be 9.58 substitutions per year, similar to the number expected based on SNP rates calculated for *S*. Typhimurium (Marzel et al., 2016; Octavia et al., 2015).

### Persistent isolates show stable plasmid and AMR gene carriage within a patient during persistence

Plasmids are a known, flexible component of the *Salmonella* genome that facilitate horizontal gene transfer and often carry antibiotic resistance genes, which could potentially contribute to a long-term persistence in an infected host (Buchwald & Blaser, 1984; Gal-Mor, 2019; Merselis et al., 1964; Murase et al., 2000). To determine the extent to which plasmid or antibiotic resistance gene (ARG) content was associated with persistence, we characterized plasmids and ARG content in each isolate using MOB-suite and RGI, respectively, and determined differences in their repertoire between early and late isolates for each patient. Overall, MOB-suite predicted 1,193 plasmids among the 639 *Salmonella* isolates (1.87 plasmids per isolate, on average), which were clustered into 96 distinct homology groups based on genomic distance estimation using Mash [**Supplemental Table 2**]. The most prevalent plasmid groups also tended to be serovar specific and widely held among members of that serovar, supporting prior work showing a serovar-specific distribution of plasmids among salmonellae including the pSLT virulence plasmid of *S*. Typhimurium, the pSEN plasmid of *S*. Enteritidis (Platt et al., 1988), and the pESI plasmid and *S*. Infantis [**Supplemental Figure 1B**] (E. Cohen et al., 2020; Nadin-Davis et al., 2020). Here, we observed a previously unreported association between the p2.3 plasmid and *S.* Enteritidis in the majority of persistent isolates of this serovar. Conversely, plasmids pSH14-028_2 and pSAN1-1677 (in red and orange, respectively) did not appear to be serovar specific, but rather were found in 133 and 93 isolates, respectively, across the many diverse serovars of *Salmonella* of the persistent isolates dataset.

Interestingly, when we compared plasmid carriage across early and late same-patient isolates using plasmid group assignments [**Supplemental Table 3**], we found plasmid content to be generally stable over time with only little evidence for plasmid gains and losses [**Supplemental Figure 1A**]. Moreover, among same-patient isolates changing in plasmid content over time, there was no association between persistence and the number of gains or losses of a plasmid group or plasmid group frequency [**Supplemental Data, Supplemental Figures 1C,D**]. Similarly, we found that ARG content of most same-patient isolates, (whether predicted to be plasmid or chromosomally-encoded), remained relatively stable [**Supplemental Figure 2D**], with only a handful of late isolates gaining large numbers of resistance genes [**Supplemental Table 4**]. Late isolates gaining ARGs were not associated with a specific serovar, plasmid, or time interval between early and late isolates.

### Persistent *Salmonella* isolates are enriched for genetic changes in global regulators

We hypothesized that some *Salmonella* adaptations enabling long-term carriage in human hosts would be conserved across serovars and patients at the gene and/or functional level. To search for such adaptations, we annotated and compared the locations of intergenic or coding variants, including SNPs, insertions, and deletions, that differed between early and late same-patient isolates across all cases. Despite 388 variable sites in CDS regions, only a small fraction (5.5% or 259 of 4,634) of genes were changed. Of these, 26 genes were found to contain variants between the early and late isolates in at least two different patients, including both synonymous and nonsynonymous mutations [**Table 1**, **Figure 2A**]. There were an additional 135 variable intergenic sites; however, only 5 intergenic regions had variants in two or more patients and the only region with variants in more than two patients was that to which the closest gene is downstream and encodes a YjiH-Family protein, HAD6495080, and is shared by 4 patients [**Supplemental Table 1**].

**Figure 2:**
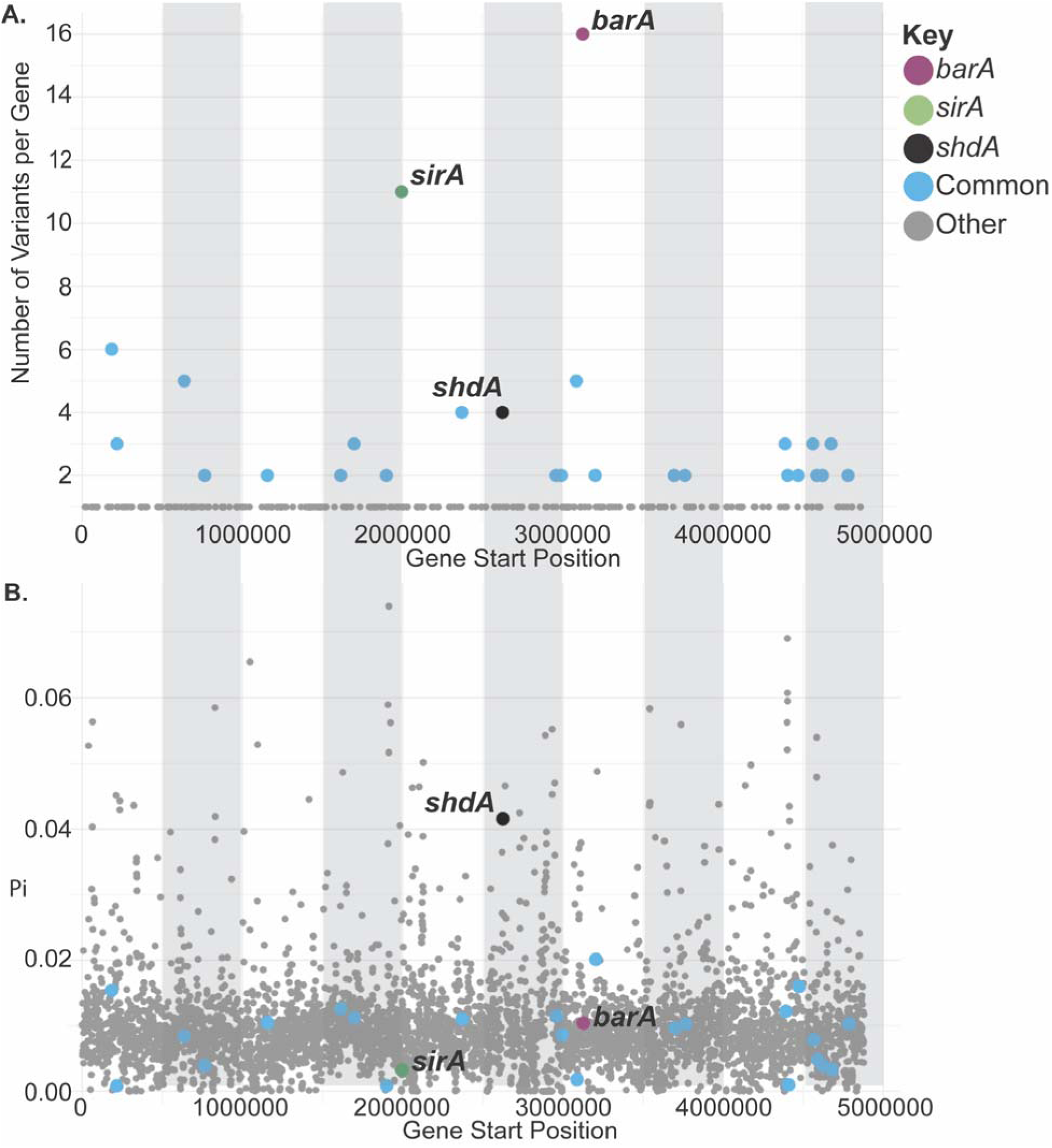
Most commonly mutated genes do not have high nucleotide diversity. **A)** The number of distinct variants between early and late same-patient isolates per gene for all genes that contain variants, plotted by gene start position. **B)** Average nucleotide diversity, □, per gene in the reference genome. *barA* in purple, *sirA* in green, *shdA* in black, common genes (genes with variants in at least 2 patients) in light blue, and all other genes in gray.

**Table 1:**
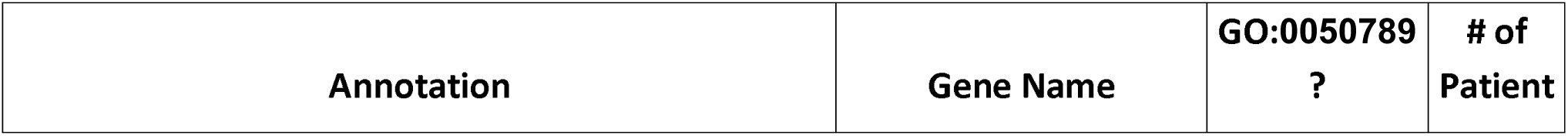
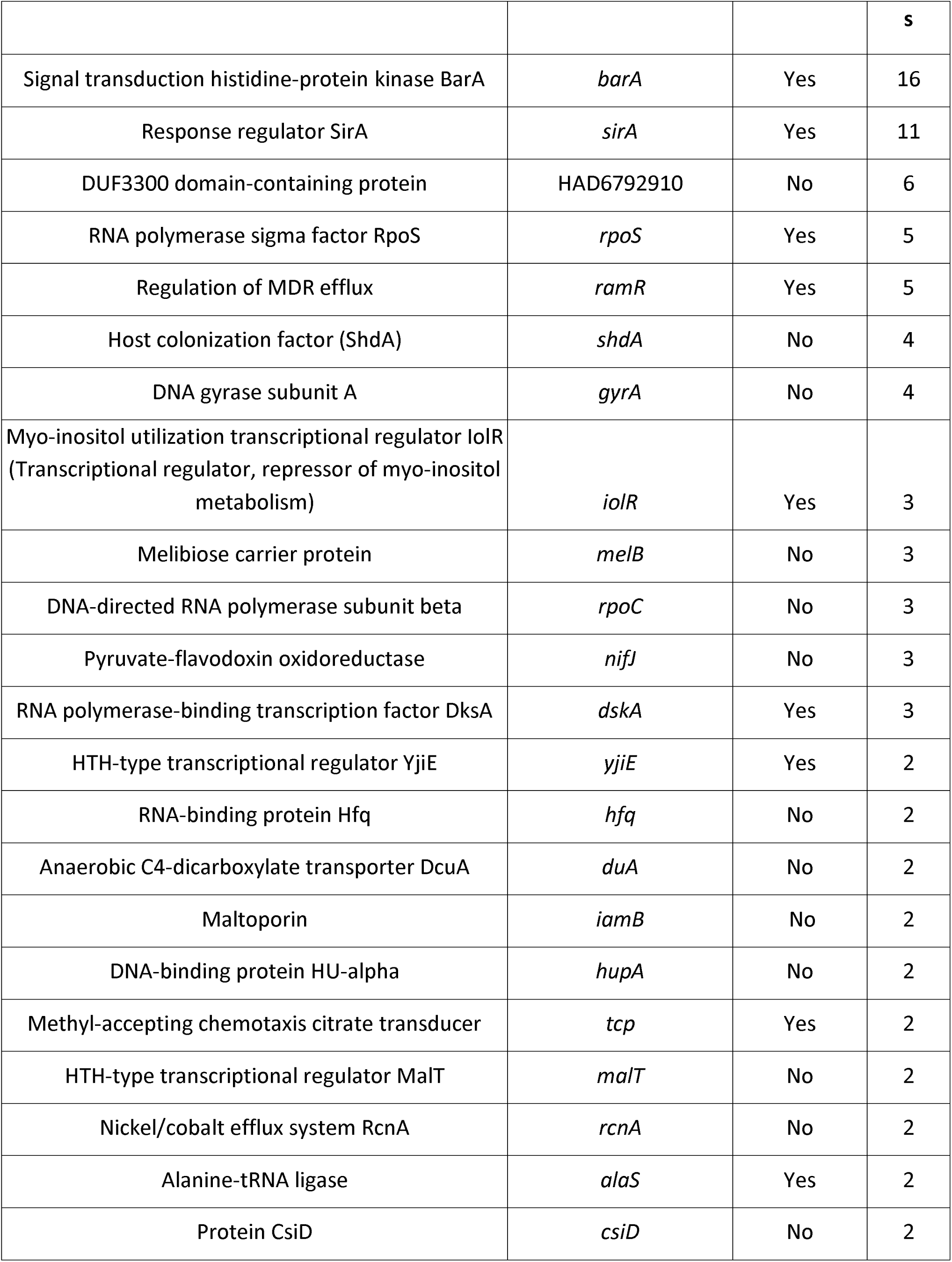
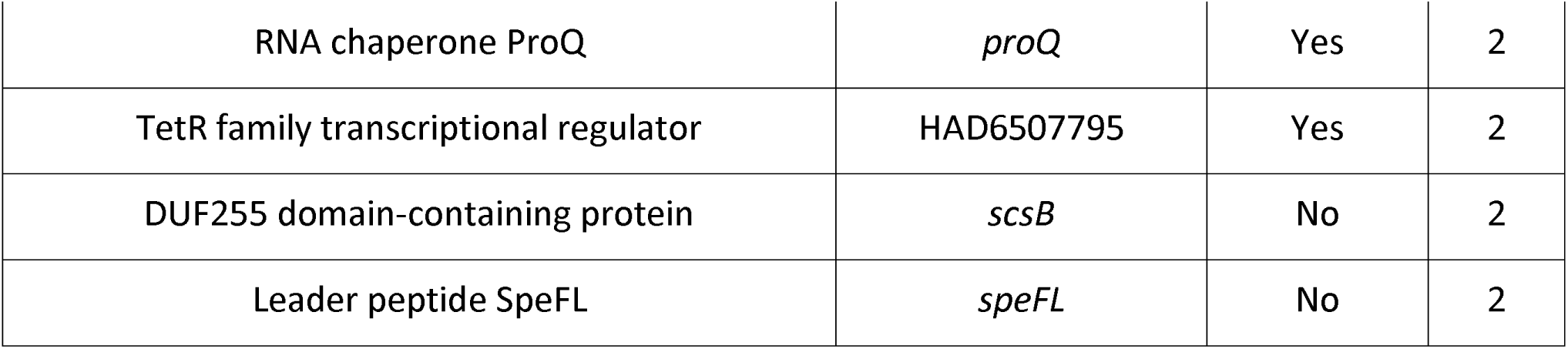
Genes with variants between early and late same-patient isolates identified in at least 2 different patients. Annotation, gene name, whether that gene is annotated with the GO term 0050789 (regulation of biological process), and the number of different patients with a variant in that gene are listed.

To determine convergence at a pathway level, we performed a GO term enrichment analysis of genes acquiring nonsynonymous or nonsense mutations found that all 22 enriched (FDR less than 0.05) Biological Process GO terms were involved in regulation and transcription, indicating a larger trend toward variation in regulatory processes during persistence [**Supplemental Table 5**]. To explore this further, we interrogated the GO term 0050789, regulation of biological process, which was enriched (FDR of 0.0140) in nonsynonymous variants arising in isolates from diverse serovars. Nearly a quarter (103 of 423) of all observed variants were in genes with this GO term, and nearly half (68 of 150) patients with at least one variant had a variant in a gene with this GO term [**Figure 1C, Supplemental Table 6**]. Of the 26 genes listed in **Table 1**, 11 were assigned GO term GO:0050789 though we noted that additional genes in **Table 1** have also been implicated in regulation in *Salmonella*, including *malT*. Therefore, we concluded that genetic variation emerging over time in these global regulatory genes was common across many diverse *Salmonella* serovars [**Figure 1C**]. Also in **Table 1** are a number of genes implicated in antibiotic resistance including *gyrA*, *tetR*, and *ramR*. A SNP in the *gyrA* gene was acquired in the late isolate during persistence in four different patients. In Patient 260, the SNP is located within the quinolone resistance-determining region (QRDR), causing a glycine to aspartate substitution at position 81, which resulted in nalidixic acid resistance in the late isolate.

While variants between same-patient early and late isolates were found at numerous genomic loci, variants in two genes, *barA* and *sirA,* were overwhelmingly the most frequent [**Table 1**, **Figure 2A**]. Strikingly, we observed an accumulation of 27 different variants in *barA* and *sirA* in 24 different patients, across 14 different *S. enterica* serovars [**Table 1**]. *Salmonella* strains in two patients gained mutations in both *barA* and *sirA*, and one patient accumulated two different *barA* mutations. In addition, though found in only a single patient, one pair of isolates had a mutation in *hilD,* one of the transcriptional regulators of SPI-1, activated by the SirA/BarA system. All mutations accumulating over time in these genes were predicted to lead to either missense or nonsense changes.

### Convergent evolution in SirA/BarA and other regulatory genes not due to overall increased mutation rate at these loci

While the higher frequency of sequence variation between early and late isolates in the genes listed in **Table 1** suggested a selection for these variations during persistence, we reasoned that it may also be due to a higher overall mutation rate in these loci. To determine if this was the case, we compared the nucleotide diversity, □, (Danecek et al., 2011) for *barA* and *sirA*, as well as the other genes in **Table 1**, to that of all other genes in the reference genome using i) all possible isolate pairs within our collected set of early *Salmonella* isolates ii) a more diverse set of 875 *S. enterica* complete genomes that were clustered at 99% kmer similarity down to a set of 177 representative genomes [**Supplemental Table 8**]. In order to generate an alignment for each gene in our reference genome, *S*. Typhimurium SL1344, we aligned reciprocal best BLAST hits between each refseq genome and our reference. For each gene’s alignment, we calculated □ [**Supplemental Figure 3**]. The average nucleotide diversity of genes calculated from either set of genomes pointed to the same conclusion. All but 2 genes were within one standard deviation of the mean nucleotide diversity of all genes in the genome [**Figure 2B**]: *shdA* (□=0.042) encoding an outer membrane fibronectin-binding protein previously shown to be highly diverse across *Salmonella* (Urrutia et al., 2014) (**Supplemental Table 9**). Taken together, these results suggest that variations in *sirA* and *barA* reflect convergent evolution of diverse serovars in multiple patients that enable *Salmonella* to overcome common barriers to chronic infection.

### BarA and SirA variants in persistent isolates lead to downregulation of virulence genes encoded in SPIs 1 and 4

We next sought to characterize the transcriptional consequences of *sirA*/*barA* sequence variation. We selected four pairs of early and late same-patient isolates separated by the fewest number of variants including either a nonsynonymous SNP or deletion in *barA* (2 pairs) or *sirA* (2 pairs) and compared their transcriptomes by RNA-Seq, following growth to the mid-exponential phase in LB. Although these pairs of isolates were separated by only 1-2 mutations (including the *barA* or *sirA* mutation), we found dozens of genes differentially expressed between each pair of isolates. Strikingly, the sets of downregulated genes were largely overlapping, with numerous genes encoded in SPI-1 and SPI-4 and genes encoding their associated effector proteins significantly downregulated in all four patients [**Figure 3, Supplemental Figure 4, Supplemental Table 7**]. Among genes upregulated between pairs of late and early isolates, *ompF*, which encodes the OmpF porin protein, was the only gene upregulated in all four patients. Of note, the two Isolate pairs with variations in *sirA* shared ten upregulated genes, including four involved in propionate degradation, *prpBCDE*. Interestingly, it has recently been shown that propionate metabolism is linked to intestinal expansion by *S*. Typhimurium in the inflamed gut (Shelton et al., 2022). Taken together, these findings suggest that independent mutations in *barA* and *sirA* acquired by late isolates in independent patients lead to largely (though not completely) overlapping transcriptomic changes, which include the downregulation of numerous SPI-1- and SPI-4-encoded factors and associated effectors.

**Figure 3:**
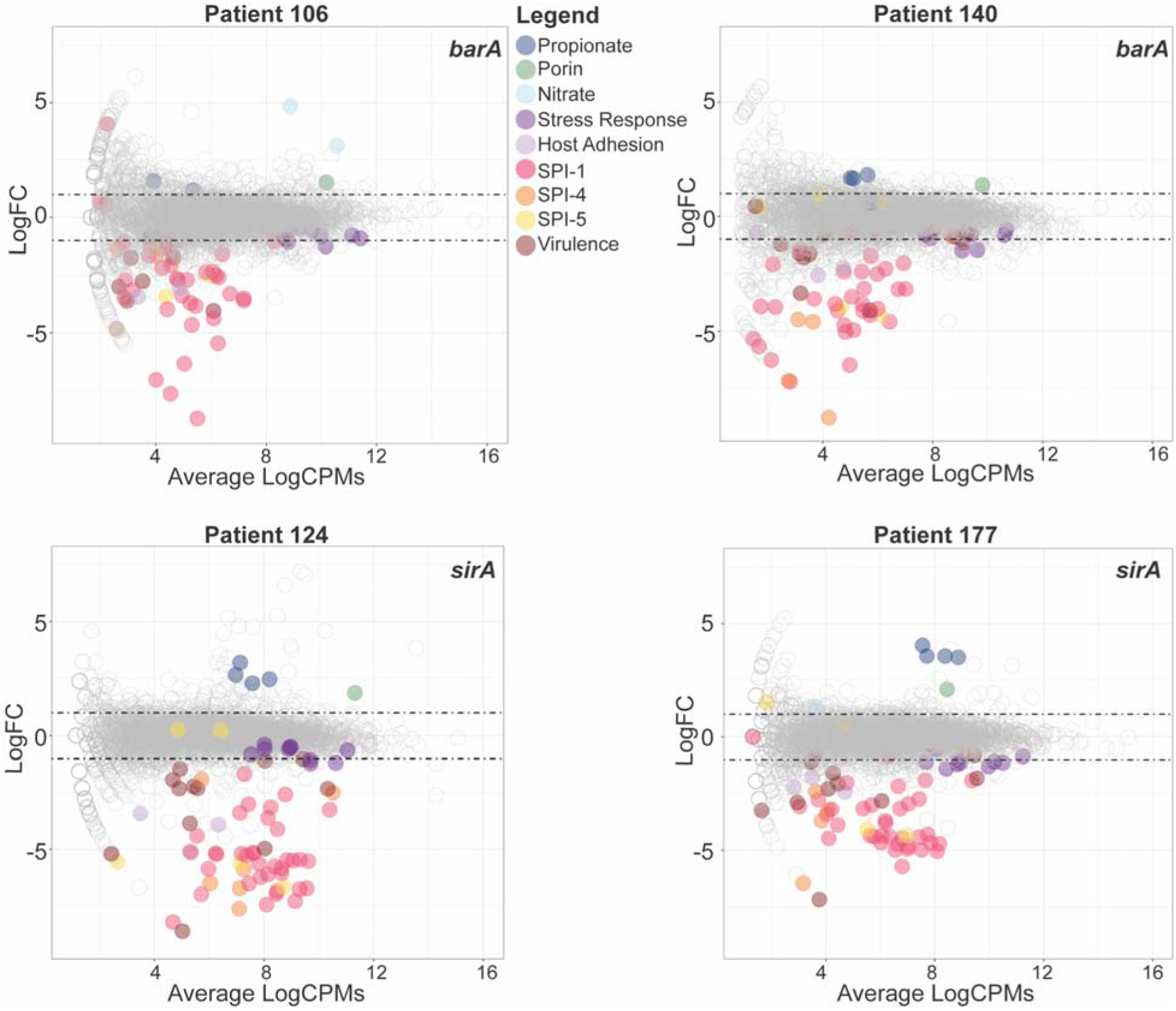
Differentially expressed *Salmonella* virulence genes in patients with *sirA/barA* variants. MA plots showing log fold-change (logFC) vs log average expression (logCPM) for four different patients, two with *barA* mutations (Patients 106 and 140) and two with *sirA* mutations (Patients 124 and 177). Genes of interest that were found to be differentially expressed in either both *barA* variant strains, both *sirA* variant strains or in all four variant strains are colored based on annotation: propionate degradation (blue), porin (green), nitrate reduction (light blue), stress response (purple), host adhesion (light purple), SPI-1 (red), SPI-4 (orange), and SPI-5 (yellow), virulence (maroon). The complete list of DEGs is shown in **Supplemental Table 7**.

### Genetic variants in *barA* and *sirA* attenuate *Salmonella* virulence in the mouse model

Because genes encoded in SPI-1 and SPI-4 are required for *Salmonella* virulence, we hypothesized that the observed downregulation of SPI-1 and SPI-4 genes mediated by sequence variations in *barA* and *sirA* would lead to a decrease in *Salmonella* pathogenicity. To test this, we performed competitive infection in streptomycin pre-treated C57BL/6 mice, an established *Salmonella* pathogenesis model (Palmer & Slauch, 2017). For each of the four pairs of isolates for which we performed RNA-Seq above, equal amounts of the differentially marked early and late same-patient isolates were used to co-infect mice. At four-days post infection (p.i.), we plated spleen, liver, cecum and colon homogenates onto antibiotic selective plates and calculated the relative bacterial loads of late vs. early isolates. For each pair, the later isolate (carrying a *barA* or *sirA* mutation) was recovered from the mice at significantly lower quantities than the early isolate in nearly all body sites, indicating that these *sirA/barA* mutations attenuate virulence in this mouse model by reducing the ability of the strain to colonize the gastrointestinal tract and in some cases systemic sites as well [**Figure 4**].

**Figure 4:**
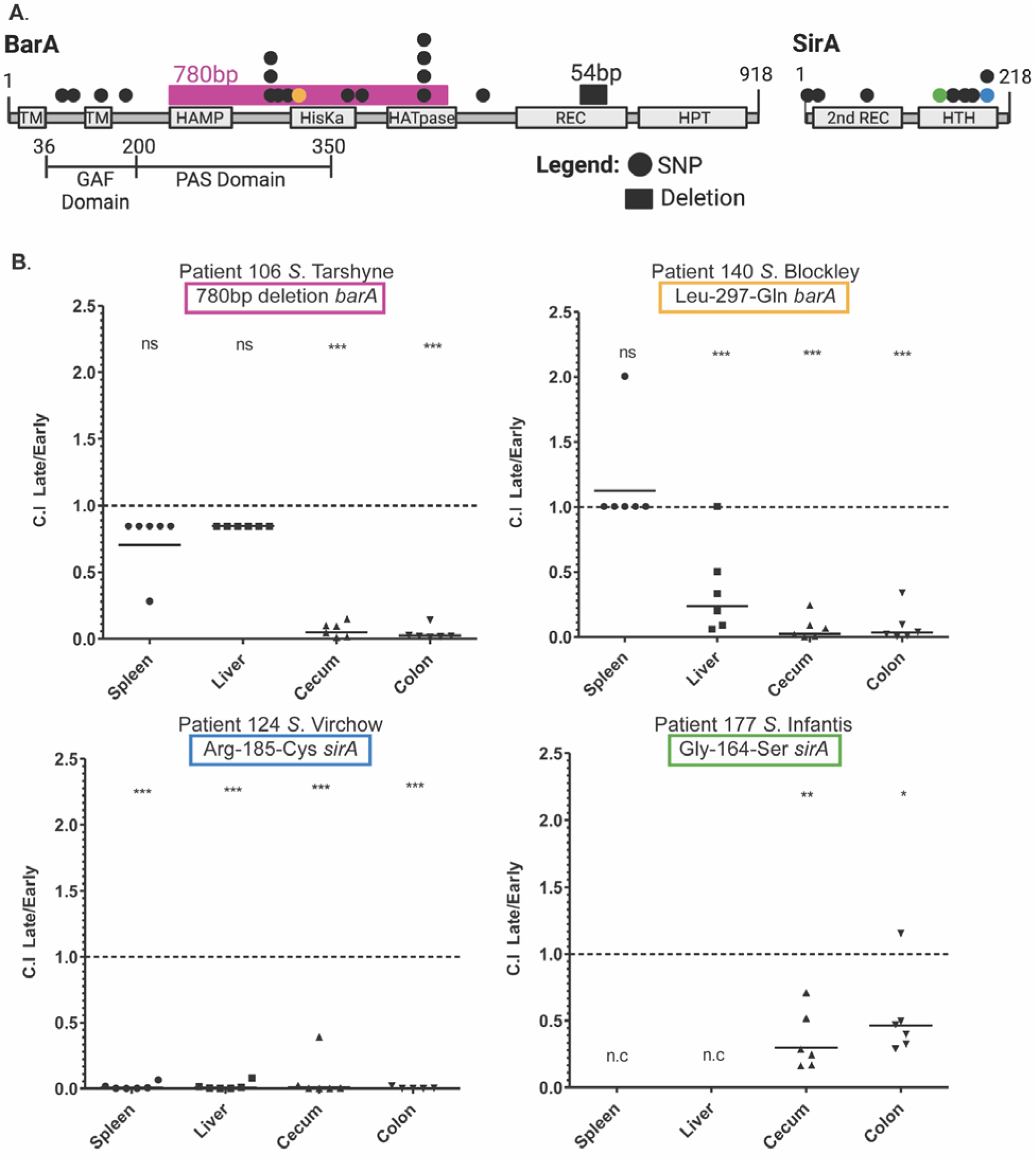
Variation in *barA* and *sirA* acquired during persistence attenuates *Salmonella* pathogenicity. **A**) Diagram of the BarA and SirA proteins with domains indicated by boxes: TM = transmembrane domain; HAMP = histidine kinase, adenyl cyclase, methyl-binding protein, phosphatase domain; HisKa = dimerization and phosphoacceptor domain; HATpase = histidine kinase-like ATPase; REC = primary receiver domain; HPT = secondary transmitter domain; 2nd REC = secondary receiver domain; HTH = helix-turn-helix domain. SNPs are indicated by circles and deletions by rectangles, the 4 colors indicate the SNPs and deletion that were tested in the mouse model. **B**) Competitive index (CI) of late vs. early isolates with variants in *barA* or *sirA* after coinfection of C57BL/6 mice. CI was calculated for the spleen, liver, cecum, and colon of the mice. ***, p<0.001, ns stands for not significant. The *Salmonella* serovar and location of the SNP in *barA* or *sirA* is shown at the top of each graph. The colored box around the location matches the color of the SNP or deletion in (A).

## DISCUSSION

Using comparative genomics across a large set of persistent *Salmonella* isolates from diverse serovars, we identified convergent within-host mutations in global regulatory genes across nearly 50% of patients, including in the genes encoding the SirA/BarA TCRS. Using RNA-Seq, we found that these mutations led to the downregulation of many genes encoded in SPI-1 and SPI-4 and the genes encoding T3SS-1 associated effector proteins. Finally, competition experiments in a mouse model of salmonellosis revealed that the late isolates harboring a mutation in either *barA* or *sirA* were significantly less virulent than the early isolates. Taken together, we propose a model by which mutations in the genes encoding the SirA/BarA TCRS lead to the downregulation of SPI-1 and SPI-4 genes, and to attenuated virulence allowing the infection to persist in the host.

### Adaptations in global regulators may be a common theme of bacterial persistence in humans

GO enrichment analysis of the nonsynonymous variants between early and late same-patient isolates revealed significant enrichment of genes involved in transcriptional regulation, or regulation of biological processes. Remarkably, despite very few mutations separating isolate pairs, variations in a global regulatory gene between early and late isolates were found in 46% of patients, including 24 patients with variants in the *sirA/barA* genes. The SirA/BarA TCRS is a well-conserved membrane-associated sensor kinase and cognate response regulator that controls *Salmonella* virulence through SPI-1.

While our findings are the first to show convergent adaptation in SirA/BarA in persistent *Salmonella*, recent studies in other bacteria that cause persistent human infections have also identified within-host adaptations in global regulatory genes, including in orthologs of SirA/BarA, GacA/S, in persistent *Pseudomonas aeruginosa* infection of the cystic fibrosis (CF) lung (Bartell et al., 2019; Marvig et al., 2015). Also in the context of CF, mutations in the genes encoding the *Staphylococcus aureus* virulence transcriptional regulators AgrA and RsbU were found to be under positive selection (Long et al., 2021) as were the global regulatory proteins PhoQ, BigR, SpoT and CpxA in *Achromobacter xylosoxidans* infection (Khademi et al., 2021). Outside of CF, mutations in the two-component regulatory systems, SsrA-SsrB and VraS/VraR, controlling aspects of *Staphylococcus aureus* pathogenicity were described in a clinical case of a persistent *S. aureus* diabetic foot infection (Lavigne et al., 2021) and this theme was also observed for persistent lung infections caused by the environmental pathogen, *Mycobacterium abscessus* (PhoR, Crp/Fnr, EngA, a TetR family member, and IdeR) (Bryant et al., 2021). Taken together, our findings and these studies suggest that positive selection of mutations in global regulatory pathways may be a common path for diverse pathogens to transition from acute to persistent infection.

### Variants in BarA and SirA lead to downregulation of virulence genes in SPI-1 and attenuate virulence of persistent isolates in the mouse model

To understand the mechanism by which mutations in key virulence regulators could lead to persistence in the human host, we profiled the transcriptional effects of mutations in the SirA/BarA two-component system carried by late isolates. Using RNA-Seq to compare the transcriptomes of early and late same-patient isolates, we found that the genes downregulated in the persistent isolates in both patients were strongly enriched for genes in SPI-1 and SPI-4. SPI-1 is known to control invasion of cells in the intestinal epithelium leading to enteritis through a T3SS that is able to inject effector proteins directly into the cytoplasm of the host cells (Lou et al., 2019). SPI-4 which encodes the invasin protein SiiE has been shown to be coregulated with SPI-1 and is believed to have a complementary role to SPI-1 during the enteric phase of *Salmonella* pathogenicity (Kiss et al., 2007). In contrast, we found far fewer shared upregulated genes in the persistent isolates, the only commonly upregulated gene being *ompF*, one of the major porins in *Salmonella*. It has been shown in *S.* Typhimurium that OmpF is recognized by macrophages, triggering a signal cascade that results in phagosomal maturation (Ipinza et al., 2014; Negm & Pistole, 1998). OmpF has also been shown to be involved in antibiotic resistance in *E. coli* and other bacteria (Choi & Lee, 2019). Therefore, it is possible that induced expression of OmpF contributes to *Salmonella* persistence.

To further assess how variations in the SirA/BarA TCRS contribute to persistence, we used the mouse model for *Salmonella* to compete early isolates against late isolates with variants in either *barA* or *sirA* for two different patients each. We found that, in all four patients, the persistent (late) isolate with the *barA* or *sirA* variant was less virulent compared to the early isolate. In all cases, the early isolate significantly outcompeted the late isolate and caused higher bacterial burden in the cecum and the colon. The Leu-297-Gln mutation in BarA in *S*. Blockley was also significantly outcompeted by the WT in the liver, while the Arg-185-Cys mutation in *sirA* in *S*. Virchow was significantly outcompeted by the WT in all four organs tested, including the spleen. Based on these results, we propose a mechanism of *Salmonella* persistence whereby mutations in global regulatory genes that result in virulence attenuation are positively selected during infection and facilitate sustainable long-term infection “under the radar” of the host immune system

### Persistent isolates show stable plasmid and antibiotic resistance gene carriage within a patient over the infection period

While antibiotic treatment was previously suggested as a potential risk factor for the development of persistent infection (Buchwald & Blaser, 1984; Gal-Mor, 2019; Lai et al., 1992; Merselis et al., 1964; Murase et al., 2000), our current study did not find a clear molecular or genomic evidence to support such an association. For most persistent cases, carriage of accessory antibiotic resistant genes did not appear to change over time, and no AMR genes were selectively gained or lost in persistent strains. Nevertheless, in a few cases we were able to identify late presence of SNPs in core genes, which confer antibiotic resistance. For example, the Gly-81-Asp SNP in the quinolone resistance-determining regions (QRDR) of *gyrA* in the late isolate of patient 260 was found to lead to a quinolone resistance phenotype (data not shown), suggesting that point mutations in the core genome during persistence might be a more frequent mechanism for antibiotic resistance than horizontal acquisition of AMR genes. Because our retrospective study focused on changes within the host during persistence, we were not able to evaluate prospectively whether isolates that will become persistent are loaded with more AMR genes upon entering the host than are isolates that do not become persistent.

Similarly, while we identified a diverse repertoire of plasmids in the collected dataset, we found that none of them were more likely to be gained or lost during persistence in the human host. We found that plasmid carriage did not increase with time/ duration of infection, nor were any particular plasmids found to be associated with persistence. These results suggest that plasmid dynamics and acquisition of antibiotic resistance are not a major driving force for *Salmonella* persistence. We found that many plasmids appear to be serovar specific, reproducing some known plasmid-serovar associations (E. Cohen et al., 2020; Nadin-Davis et al., 2020) as well as finding potential new associations. The stable presence of plasmids during long-term infections in humans suggests that their maintenance does not pose a significant metabolic burden during persistent infection. In addition, this could also indicate that the site of persistence is not enriched with high diversity microflora such as the gut lumen that permit horizontal gene transfer (Lerner et al., 2017), but rather an isolated, possibly intracellular, site such as the gallbladder, lymph nodes, or hemophagocytic macrophages.

### Additional considerations and future studies

While “early” isolates in our dataset were taken upon first culture-confirmed *Salmonella* diagnosis, we can not be certain that all early samples were taken at the initial onset of disease. Indeed, the presence in the blood of some “early” isolates may be indicative of the fact that these were taken at a later stage of infection. Because of this, we may have missed some transitions to persistence within patients. This may be why, in some patients, we do not find any variants between the early and late isolates whatsoever. To this point, we did observe several patients with presumably inactivating mutations (insertions, deletions, or nonsense mutations) in *barA* or *sirA* shared by the early and late isolates, which may point to a transition to persistence having already occurred. Additionally, we chose to use a reference genome against which to compare all isolate genomes, because we were interested in finding a trend that occurred across the phylogeny and that was serovar-independent. While this method proved to be fruitful, we could be missing additional strain or serovar-specific adaptations to a persistent life-style that are in genomic regions not present in the reference *S*. Typhimurium genome. Lastly, because we were only able to perform RNA-Seq on a limited number of isolates, we chose to focus on the *barA* mutations that were shown to have an effect in the *in vivo* mouse model. Future work could include comparative transcriptomics across all early and late same-patient isolate pairs, which may reveal convergent transcriptional pathways resulting from diverse genetic variants. Future work to understand how the host immune system responds to the decrease in virulence would also be illuminating.

While unanswered questions remain about how bacterial pathogens establish and maintain persistent infections, this study provides key insights into the genetic adaptations underlying *Salmonella* persistence and the transcriptional mechanism by which they influence virulence. Not only does our large set of whole genome sequences of persistent NTS of *Salmonella* provide an invaluable resource for the community, our study also serves as a template for how mechanisms of persistence can be studied in other bacteria. The size and scope of our study allowed us to pinpoint convergent evolution not only at the nucleotide and gene level, but also at the pathway level where mutations are acquired in different genes that all share a common function. The discovery of these conserved pathways to persistence makes the discovery of potential therapeutics to block or treat persistence all the more possible.

## MATERIALS AND METHODS

### Sample collection and isolation

From one of the largest retrospective studies of *Salmonella* persistence including 48,345 culture-confirmed salmonellosis cases that occurred in Israel over 17 years (between 1995 and 2012) (Marzel et al., 2016), we assembled a collection of 639 longitudinal *S. enterica* isolates representing 49 NTS serovars obtained during persistent infections from 256 patients. Each patient was represented by 2-5 independent isolates obtained during different stages of infection that lasted between 30 and 2,001 days from the first culture-confirmed diagnosis. Isolates were plated on XLD agar plate and a single colony was grown in LB for O.N. at 37°C under aerobic conditions on a Roller Drum. The next day the DNA was extracted from 500 µl of culture using the GeneElute bacterial genomic DNA kit (Sigma-Aldrich). gDNA was quantified, assessed for its quality and whole genome sequenced.

### Whole genome sequencing and variant analysis

Purified DNA (0.2 ng) was used as input into a miniaturized version of the Nextera-XT Library Preparation Kit (Illumina Inc.). All reactions were scaled to one-fourth their original volumes. Libraries were constructed according to the manufacturer’s instructions with the following minor modifications to boost the insert size, thereby reducing read overlap during paired-end sequencing: i) tagmentation time was reduced from 5 minutes to 1 minute; individual sample libraries were pooled at equimolar concentration. The final library pool underwent an additional size selection using a 0.7X SPRI using AMPure XP beads (Beckman Coulter) and was sequenced on a NovaSeq SP at 300 cycles for 2×150 pair-end reads. Genome coverage was approximately 150-fold, on average. Resulting fastq files were checked for quality using FastQC, and checked for contamination using Centrifuge (Kim et al., 2016). In total, eight isolates were removed due to non-*Salmonella* genomic content.

BWA mem (version 0.7.12) (Li, 2013) was used to align each of the 639 genome sequences to the *S*. Typhimurium SL1344 reference genome (accession number NC_016810.1), and Pilon (version 1.12) was used to call variants between early and late isolates (Walker et al., 2014). VCF files were filtered for variants containing the filter status PASS, and variants labeled as imprecise were removed. VCF files for same-patient isolates were then compared to find all variants that differed between early and late isolates. Variants between same-patient isolates and different-patient isolates were quantified, and a cutoff of 36 bp was chosen to distinguish a persistent infection from a potential reinfection based on the longest time of infection in our dataset and previously published SNP rates (Marzel et al., 2016). Using a SNP-based alignment, Fasttree (version 2.1.14) was used to construct the phylogenetic tree of *Salmonella* isolates (Price et al., 2010). Tree visualization was performed using itol (version 6.5.6) (Letunic & Bork, 2021).

### Construction of a core genome and estimation of *in vivo* mutation rate

To determine an *in vivo* mutation rate, we first filtered sites along the *S*. Typhimurium SL1344 reference alignment that were i) low coverage (<5 reads at a position for any sample) or ii) potentially recombined based on analysis with ClonalFrameML (version 1.12) (Didelot & Wilson, 2015). From this core alignment, the mutation rate was estimated from the number of SNPs between the first and last isolate for each patient (ignoring intermediate isolates for this analysis), divided by the number of days between the two isolates. We estimated the rate and the standard deviation using the *fitdistr* function in the “MASS” package, using the Poisson distribution. Only isolate pairs that had less than 36 SNPs were included in this analysis. Lambda was found to be 0.02627, with a standard deviation of 0.02664, equating to 9.589 SNPs per year.

### Genome assembly, plasmid analysis, and antibiotic resistance analysis

Genomes were assembled using SPADES (version 3.1.1) on paired-end fastq files, using default settings (Bankevich et al., 2012). GAEMR (version 1.0.1) (http://software.broadinstitute.org/software/gaemr/) was used to check for quality of assembly and to generate standard assembly files. Draft genome assemblies were annotated with VESPER.

MOB-suite was used on the WGS assemblies to predict potential plasmids and then to cluster them by similarity, reconstruct the plasmid sequences, and make *in silico* predictions of the replicon family, relaxase type, and transferability (Robertson et al., 2020; Robertson & Nash, 2018). The Resistance Gene Identifier (RG) was used to identify potential antibiotic resistance genes in the genome assemblies using the Comprehensive Antibiotic Resistance Database (CARD) (Alcock et al., 2019).

### Nucleotide diversity analysis

We calculated nucleotide diversity, π, for each gene in the genome using two different datasets. Using the early isolate genomes in our dataset, we calculated π using the SNP calls we determined using Pilon. We used the --site-pi function in VCFtools to measure nucleotide diversity on a per-site basis across all genomes (Danecek et al., 2011). For each gene in the reference genome, we calculated nucleotide diversity per gene by taking the average site diversity divided by the length of the gene.

To calculate π using a broader database of *S. enterica* Refseq complete genomes, we used the Refseq database construction tool from StrainGE to build a database (van Dijk et al., 2022), starting with all 875 *S. enterica* Refseq complete genomes and clustering using a 90% Jaccard similarity threshold (which corresponds to 99.8% ANI) to remove highly similar genomes. This resulted in a database of 174 RefSeq genomes, representing the diversity of *S*. *enterica* [**Supplemental Table 8**]. In order to generate an alignment for each gene in our reference genome, *S*. Typhimurium SL1344, we first identified reciprocal best BLAST (RBH) hits between each Refseq genome and our reference, and then used MUSCLE (Madeira et al., 2019). We calculated □ for each gene alignment using MEGA X (Kumar et al., 2018).

### *In vivo* competition assays in mouse model

Female C57BL/6 mice (Envigo, Israel) were infected at an age of 8-10 weeks. Streptomycin (20 mg per mouse) was given by oral gavage in saline 24 h prior to infection. Mice infected with ∼4×10□ bacteria as a mixed (1:1) inoculum containing an early isolate genotype strain resistant to ampicillin (harboring pWSK29) and late isolate genotype strain resistant to kanamycin (harboring pWSK129). A mixed inoculum of *S*. Typhimurium SL1344 strains carrying pWSK29 or pWSK129 was used as a positive control. This control demonstrated a CI value of 1 which indicates similar fitness in mice.

Four days post infection, mice were humanely euthanized, and tissues were harvested aseptically for bacterial enumeration. Tissues were collected on ice and homogenized in 700 μl of saline using the BeadBlaster24 microtube homogenizer (Benchmark scientific). Serial dilutions of the homogenates were plated on XLD agar plates supplemented with ampicillin or kanamycin, incubated overnight and counted to calculate bacterial tissue burdens. The competitive index was calculated as [late isolate/early isolate]output/ [late isolate/early isolate]input.

### RNA extraction of early and persistent isolates with BarA mutations

Four patients with different *sirA/barA* mutations in the late isolate were chosen for comparative RNA-Seq analysis. Isolates were chosen due to their high genomic similarity between early and late same-patient isolates. Patients 106 and 140 had only the single *barA* deletion or SNP respectively between the early and late isolates, while patient 124 had the *sirA* SNP and one other synonymous SNP in the hypothetical protein SL1344_RS10945, and patient 177 had the *sirA* SNP and one other intergenic SNP 131 bp upstream of SL1344_RS12695, transaldolase A. Isolates were grown overnight at 37°C. The next day the bacteria were subcultured (1:100 dilution) for 3 hr at 37°C. RNA was extracted from 500 µl cultures using the Qiagen RNA protect Bacteria Reagent and RNeasy mini kit according to the manufacturer’s instructions, including an on-column DNase digest using the RNase free DNase set (Qiagen). The quantity and quality of the extracted RNA were determined by Nanodrop 2000c (Thermo Fisher Scientific). To diminish any genomic DNA contamination, RNA was re-treated with an RNase-free DNase (QIAGEN).

Cell pellets resuspended in 0.5 mL Trizol reagent (ThermoFisher Scientific) were transferred to 2mL FastPrep tubes (MP Biomedicals) containing 0.1 mm Zirconia/Silica beads (BioSpec Products) and bead beaten for 90 seconds at 10 m/sec speed using the FastPrep-24 5G (MP Biomedicals). After addition of 200 ul chloroform, each sample tube was mixed thoroughly by inversion, incubated for 3 minutes at room temperature, and spun down 15 minutes at 4°C. The aqueous phase was mixed with an equal volume of 100% ethanol, transferred to a Direct-zol spin plate (Zymo Research), and RNA was extracted according to the Direct-zol protocol (Zymo Research).

### Generation of RNA-Seq data

Illumina cDNA libraries were generated using a modified version of the RNAtag-seq protocol (Bhattacharyya et al., 2019; Shishkin et al., 2015). Briefly, 0.5-1 μg of total RNA was fragmented, depleted of genomic DNA, dephosphorylated, and ligated to DNA adapters carrying 5’-AN_8_-3’ barcodes of known sequence with a 5’ phosphate and a 3’ blocking group. Barcoded RNAs were pooled and depleted of rRNA using the RiboZero rRNA depletion kit (Illumina). Pools of barcoded RNAs were converted to Illumina cDNA libraries in 2 main steps: (i) reverse transcription of the RNA using a primer designed to the constant region of the barcoded adaptor with addition of an adapter to the 3’ end of the cDNA by template switching using SMARTScribe (Clontech) as described (Zhu et al., 2001); (ii) PCR amplification using primers whose 5’ ends target the constant regions of the 3’ or 5’ adaptors and whose 3’ ends contain the full Illumina P5 or P7 sequences. cDNA libraries were sequenced to generate paired end reads.

### Analysis of RNA-Seq data

Sequencing reads from each sample in a pool were demultiplexed based on their associated barcode sequence using custom scripts (https://github.com/broadinstitute/split_merge_pl). Up to 1 mismatch in the barcode was allowed, provided it did not make assignment of the read to a different barcode possible. Barcode sequences were removed from the first read, as were terminal G’s from the second read, that may have been added by SMARTScribe during template switching.

Reads were aligned to *S.* Typhimurium *str*. SL1344, NC_016810.1, using BWA (Li & Durbin, 2009) and read counts were assigned to genes and other genomic features using custom scripts (https://github.com/broadinstitute/BactRNASeqCount). Read counts were used for differential expression analysis, conducted with edgeR (Version 3.32.1) (Robinson et al., 2010). Differentially expressed genes were determined between the early and late isolates for each patient. Genes were determined as differentially expressed using a threshold of *p* <0.05 and a false-discovery rate (FDR) of 5%, standard settings in EdgeR. EdgeR was used to calculate CPMs, Counts Per Million, for each gene, which were used for generating heatmaps with Pretty Heatmaps (Version 1.0.12).

## Data Availability

All sequencing files from this study are available at the Sequence Read Archive [PRJNA847966].

## ACKNOWLEDGEMENTS

We acknowledge the Broad’s Microbial ‘Omics Core for generating the RNA-Seq libraries and conducting preliminary analysis for all RNA-Seq data. We also acknowledge the Broad’s Genomics Platform for sequencing. The work at the Gal-Mor laboratory was supported by grants: I-41-416.6-2018 from the German-Israeli Foundation for Scientific Research and Development (GIF); A128055 from the Research Cooperation Lower Saxony – Israel (The Volkswagen Foundation); and 2616/18 from the joint Israel Science Foundation (ISF)-Broad Institute program. The work at the Broad was supported by grants: 2616/18 from the joint ISF-Broad Institute program; U19AI110818 from the National Institute of Allergy and Infectious Diseases, National Institutes of Health, Department of Health and Human Services; and the Jane Coffin Childs Memorial Fund for Medical Research postdoctoral fellowship to A.G.. The funders had no role in study design, data collection, and interpretation, or the decision to submit the work for publication.

## SUPPLEMENTAL FIGURES

**Supplemental Figure 1:**
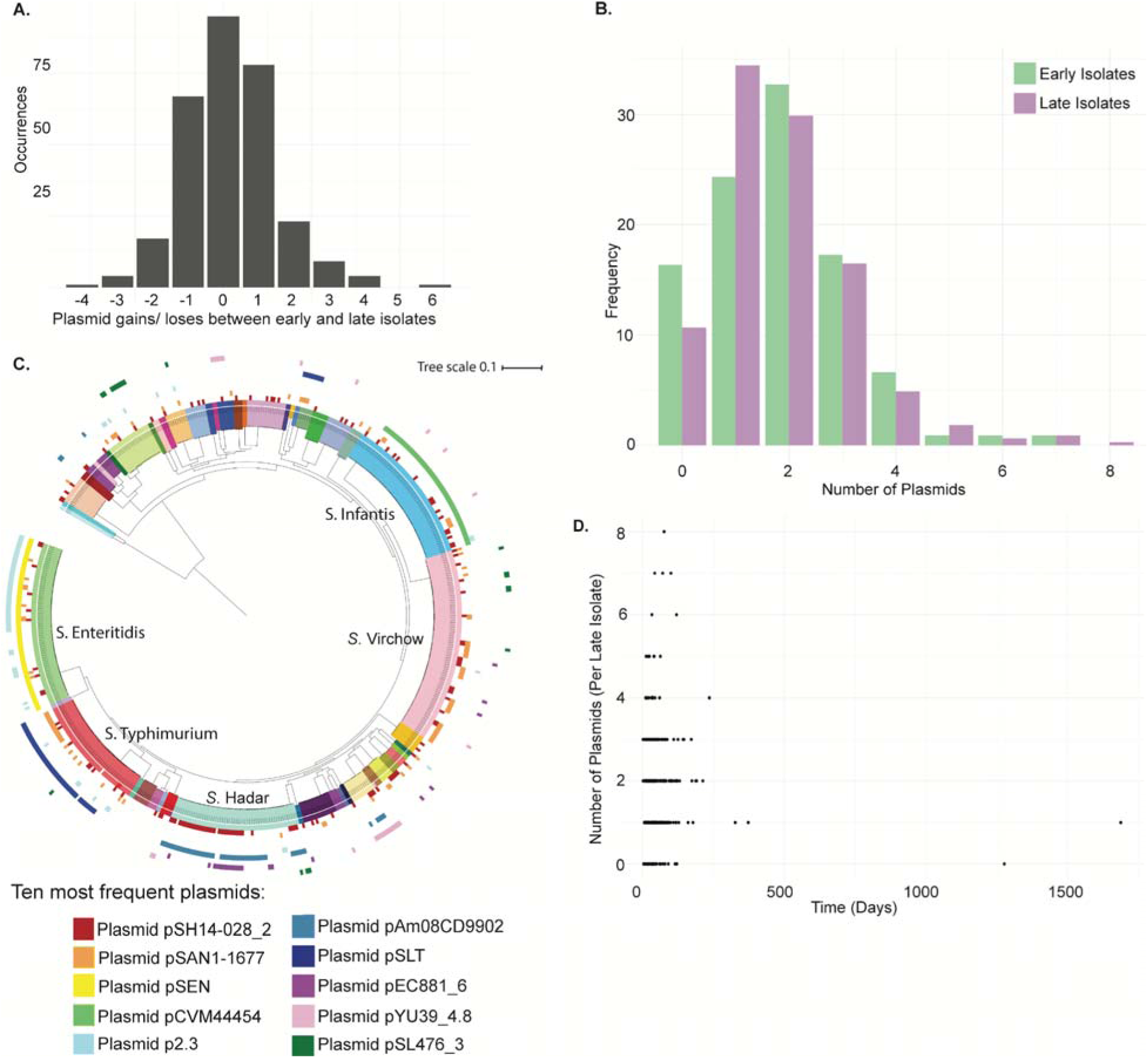
Persistent *Salmonella* has a diverse plasmid repertoire. **A)** Bar graph of the number of identified plasmid gains or losses between early and late isolates. **B)** The 10 most prevalent plasmids predicted in the persistent salmonella collection, plotted in the outer rings, with serovar indicated by the inner colored ring. **C)** The frequency at which plasmids occur in early isolates (green) and late isolates (purple). **D)** Plasmid carriage per isolate of all late isolates over time in Days.

**Supplemental Figure 2:**
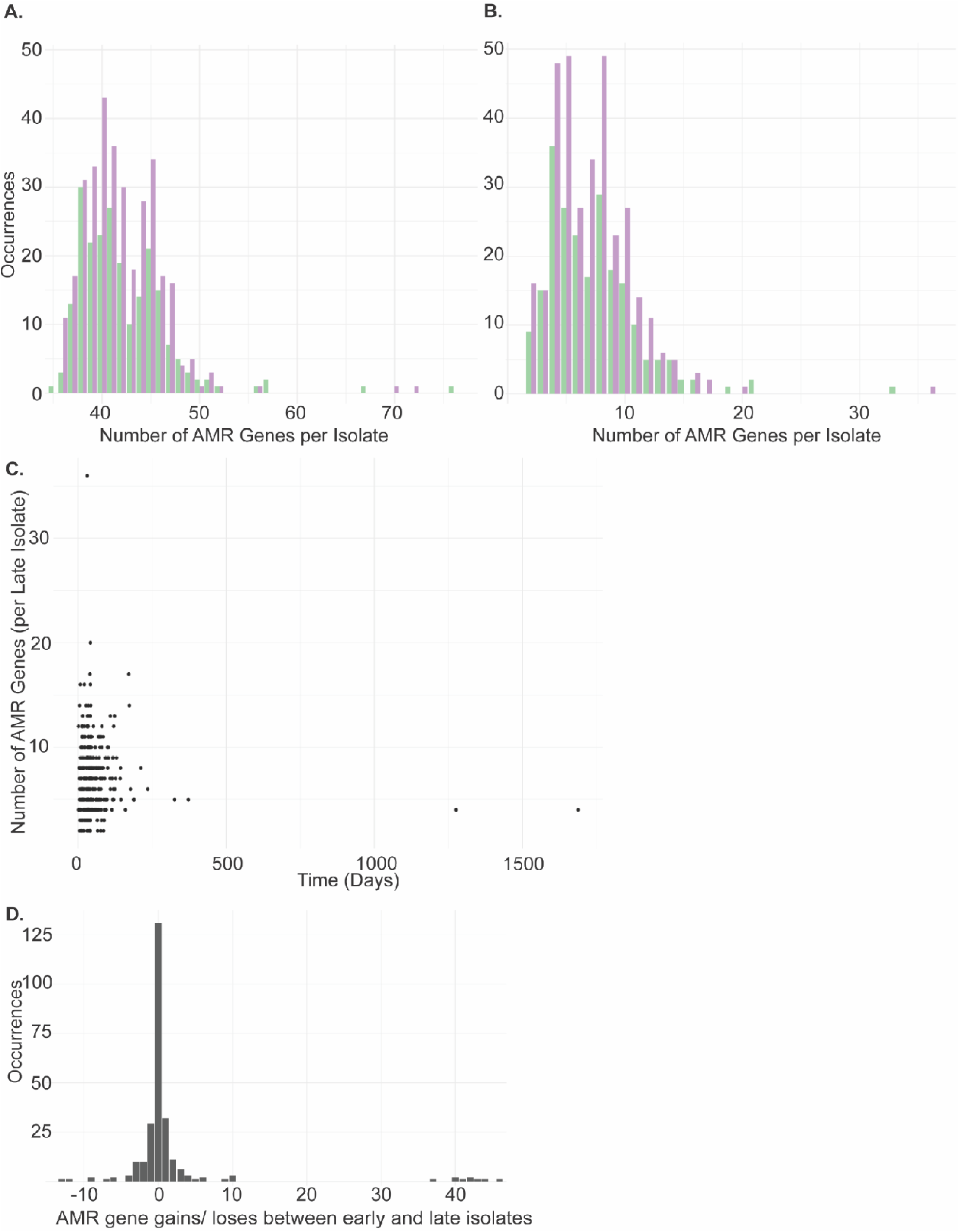
AMR gene carriage in early and late isolates as identified using RGI. **A)** The frequency at which AMR genes occur in early isolates (green) and late isolates (purple). **B)** The frequency at which AMR genes of interest occur in early isolates (green) and late isolates (purple). **C)** AMR gene carriage per isolate of all late isolates over time in Days. **D)** Bar graph of the number of antimicrobial resistance gene gains or losses between early and late isolates.

**Supplemental Figure 3:**
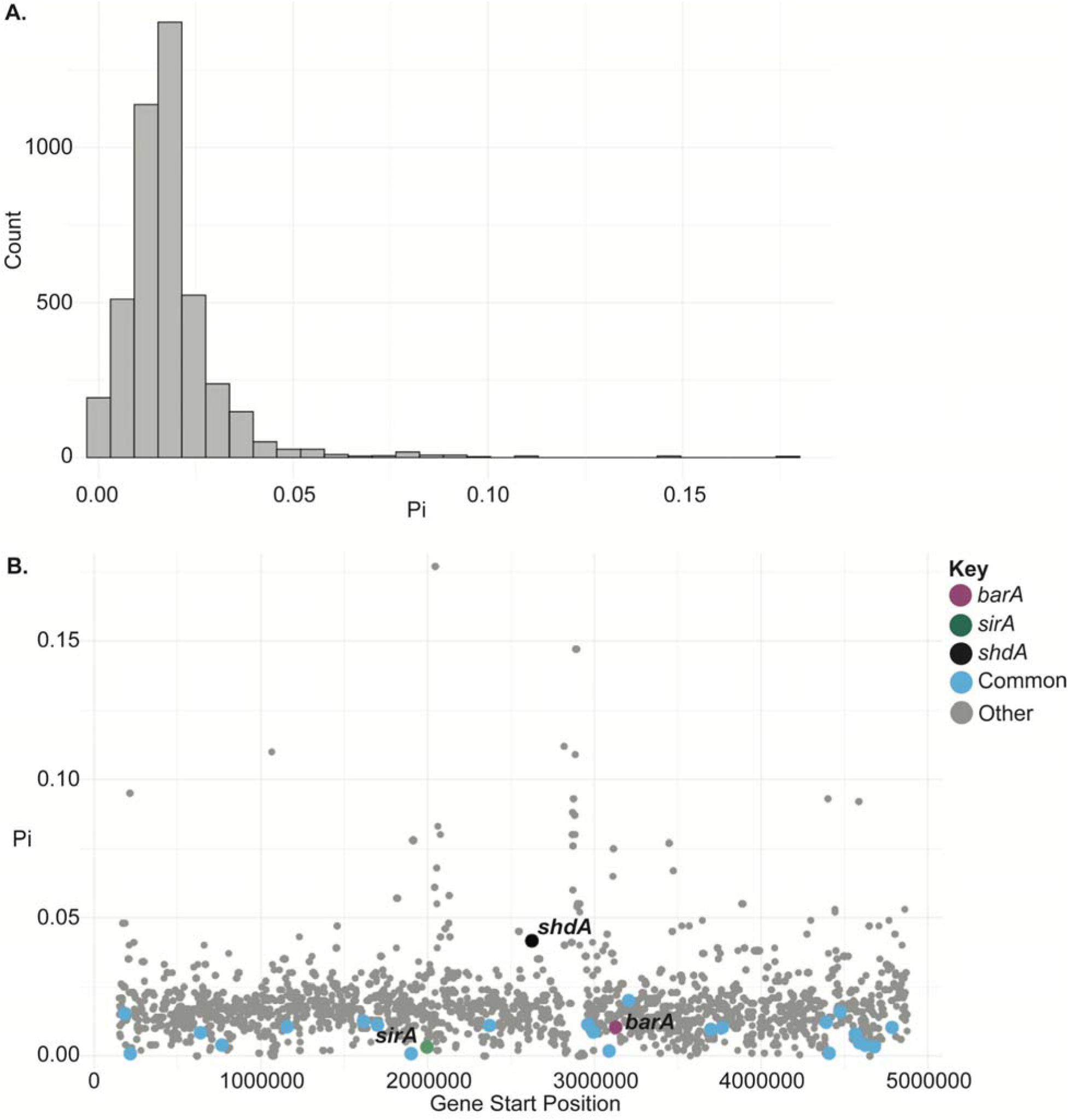
Nucleotide diversity by gene across the *Salmonella* chromosome. **A**) Histogram of nucleotide diversity for all genes in the *S.* Typhimurium SL1344 reference genome, calculated using gene alignments with 177 RefSeq genomes. **B**) Average nucleotide diversity,□, for each gene in the reference genome, plotted according to its position on the chromosome. Genes with the GO term 0050789, Regulation of Biological Process, are plotted in light blue, *barA* in purple, *sirA* in green, and *hilD* in peach. All other genes are shown in gray.

**Supplemental Figure 4:**
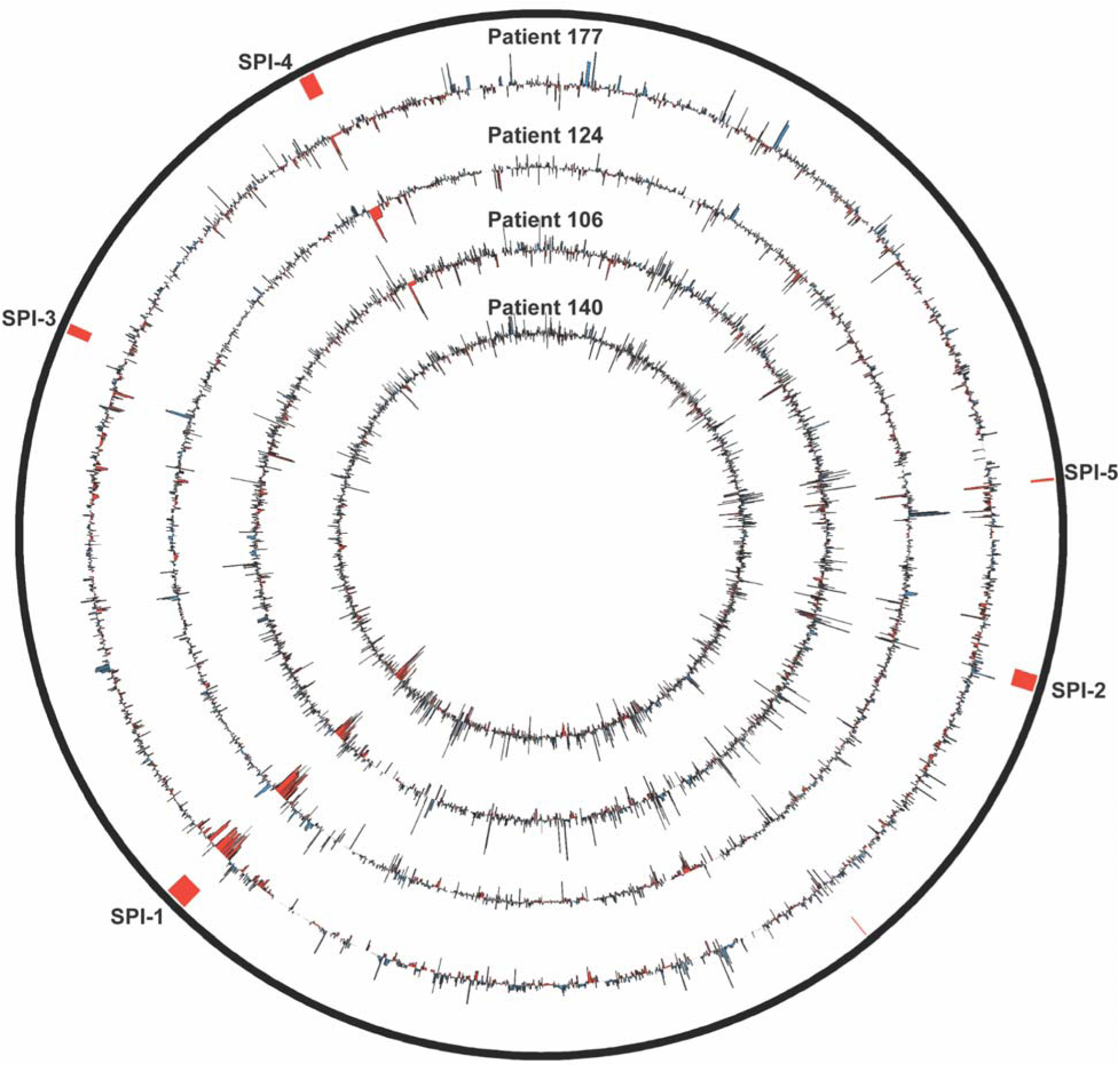
Circos plot of logFC RNA-Seq expression. The logFC for each of the four patients is indicated (patients 177, 124, 140, and 106 from exterior to interior). Blue indicates upregulated genes (positive logFC) and red indicates downregulated genes (negative logFC). SPI-1, SPI-4, and SPI-5 are indicated in the outer ring.

## SUPPLEMENTAL DATA

### RESULTS

#### Persistent isolates show stable plasmid carriage within a patient over the infection period

First, we looked at the carriage of different plasmid groups between early and late isolates to determine if there were any plasmid groups more commonly found in persistent isolates as compared to early isolates [**Supplemental Table 3**]. We found that plasmids were generally associated with specific serovars and were relatively stable between same-patient isolates. We also assessed plasmid gains and losses between same-patient isolates to determine if any plasmids appeared to be preferentially gained or lost. We determined that none of the plasmid groups predicted appeared to be gained or lost more often in early or late isolates [**Supplemental Table 1, Columns V,W**]. We next assessed the frequency or number of predicted plasmids carried by early and late isolates and found that the frequency of plasmids found by MOBsuite are very similar in early and late isolates [**Supplemental Figure 1B**]. We also looked at whether or not plasmid carriage increased as a function of time. For this analysis, we plotted the number of plasmids as a function of time (in days since last isolate sampled), ignoring the early, or time zero, isolates [**Supplemental Figure 1C**]. We found, consistent with our other analyses, that plasmid carriage appears to be relatively stable during infection and does not increase with longer infection times.

#### Antibiotic resistance gene cargo do not appear to increase with time during persistence

While we did not find any plasmids that appeared to be associated with persistence in our dataset, we wanted to look directly at antibiotic resistance genes, either chromosomal or plasmid-borne, to assess the role of antibiotic resistance gene acquisition in persistence. The emergence of antibiotic resistance has been shown to be associated with persistence in *Salmonella* (Buchwald & Blaser, 1984; Gal-Mor, 2019; Merselis et al., 1964; Murase et al., 2000); however, the connection or the involved mechanism is poorly understood. Using RGI, we first identified antibiotic resistance genes in each isolate using a read-based approach. We then assessed the frequency of predicted AMR genes in both early and late isolates, finding them to be highly similar, with an average of 42.2 and 42.0 AMR resistance genes in early and late isolates, respectively [**Supplemental Figure 2**]. We focused on a smaller group of AMR genes, known to be involved in antibiotic resistance in *Salmonella* [**Supplemental Table 10**], and assessed the relative frequency of those AMR genes between early and late isolates, and again found highly similar frequencies between early and late isolates, with an average of 7.23 and 7.11 AMR genes respectively. We also assessed whether resistance gene carriage increased over the duration of infection in late isolates and found that AMR genes do not appear to increase (or decrease) with time [**Supplemental Figure 2**]. Thus our results indicate that resistance gene repertoires in same-patient isolates appear to be stable over time, even over very long periods (∼200 days) AMR gene carriage, like plasmid carriage, appears to be relatively stable with patients during persistent infection.

